# The Interactome of DUX4 Reveals Multiple Activation Pathways

**DOI:** 10.1101/2023.09.12.557395

**Authors:** Moriya Slavin, Keren Zohar, Clothilde Claus, Dina Schneidman-Duhovny, Michal Linial, Frédérique Coppée, Nir Kalisman

## Abstract

The ectopic expression of the DUX4 protein in muscle cells is the underlying cause of Facioscapulohumeral Muscular Dystrophy (FSHD). DUX4 is a potent transcription factor that activates a large number of genes in a dysregulated manner, but the direct protein interactions involved in this activation are only partially known. Here, we tailored an affinity purification and mass spectrometry (AP-MS) analysis to the unique features and functions of DUX4, to provide a more complete view of its interactome. We also obtained and verified all-atom models for two of the major interactions by employing cross-linking and mass spectrometry (CL-MS), computational modeling, and guided mutation studies. We find that DUX4 interacts strongly with two homologous transcription activators, PTOV1 and MED25, in addition to the previously characterized CBP/p300. The interaction with the PTOV1/MED25 domain involves the wrapping of the last thirty residues of the DUX4 activation region around the domain in a very extensive interface. Hence, DUX4 has the capacity to both open the chromatin and directly recruit the Mediator complex. DUX4 also binds to all members of the RFPL4 family, which are among the strongest genes it activates. These interactions are mediated through a hitherto unrecognized motif in the DUX4 disordered linker region. This feedback mechanism suggests that DUX4 may be inhibited by its own activation products, and explains its typical pulsed expression profile. We also found SIX1 and the AP-2 complex as strong DUX4 C-terminal interactors. A separate analysis of interactions involving the N-terminal of DUX4 revealed enrichment of proteins that are involved in DNA repair following double-strand breaks. Overall, these findings reveal new activation pathways for DUX4, which may be modulated in future strategies to control its toxicity. This study also showcases the synergy between CL-MS and deep-learning based modeling for the structural elucidation of challenging protein-protein interactions.

## Introduction

Facioscapulohumeral muscular dystrophy (FSHD) is a genetic disorder characterized by progressive wasting of muscle tissue. The underlying cause is insufficient epigenetic repression of the *DUX4* gene that enables the ectopic expression of the DUX4 protein^1,2^. DUX4 is a potent transcription factor that can activate the expression of a large number of genes^3^. The dysregulated gene expression program leads to multiple cellular anomalies that include, among others, activation of cell death pathways^4–8^, oxidative stress^9–12^, morphological defects^13^, impaired nonsense-mediated decay^14^, disrupted RNA splicing^6^, and accumulation of hyaluronic acid^15^. These irregularities were suggested to have a negative impact on the regeneration of the muscle tissue^16,17^. In recent years, additional roles of DUX4 have been uncovered both in normal human development and in other diseases. In development, the early cleavage-stage embryo expresses a short pulse of DUX4 that activates an array of genes to promote the pluripotent state^18,19^. In disease, reactivation of DUX4 expression in certain types of cancers was shown to be linked to immune evasion and poor outcomes^20^, and to be a master transcriptional regulator in herpes simplex virus 1 infection^21^.

The functional regions along the sequence of DUX4 are clearly segregated (Fig. 1A). The N-terminal harbors two homeodomains that bind a specific DNA motif with high affinity^22^. The activation region at the C-terminal recruits transcription activators that facilitate the expression of nearby genes. Importantly, the deletion of the activation region abrogates the ability of DUX4 to activate genes^3^ and negates its toxic effects^23,24^. The protein termini are connected by an inherently disordered region with unknown effect on the toxicity of DUX4.

**Figure 1.**
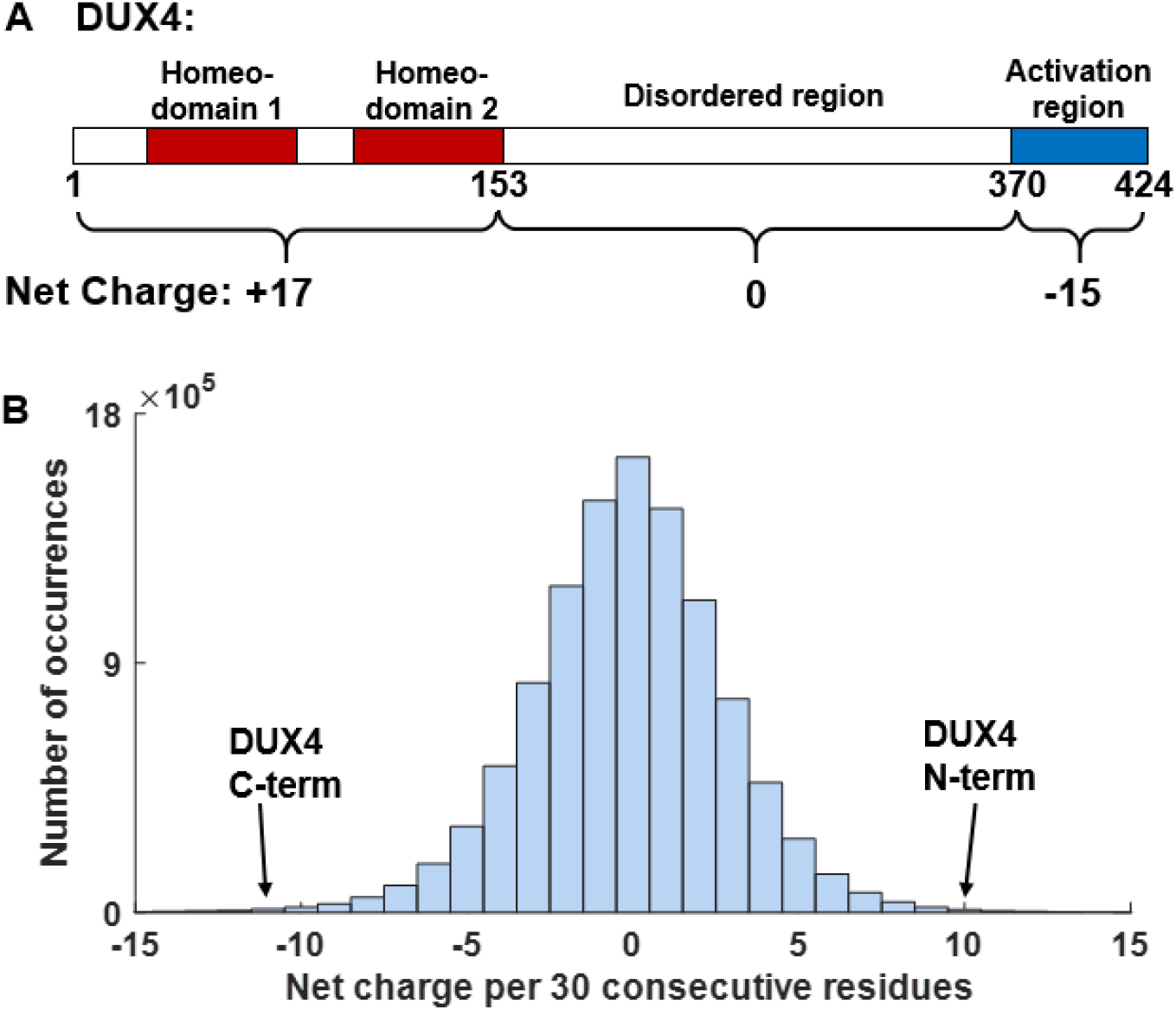
The DUX4 protein has a very polarized charge distribution. **(A)** The net charge of the N-terminal region is highly positive, while the C-terminal region is highly negative. **(B)** Histogram of the net charge per 30 consecutive residues along all the proteins in the human proteome. The values observed for the two termini of DUX4 are among the most extreme to be found in any human protein.

In order to trace the early events of toxicity, several studies have attempted to characterize the proteins that interact with DUX4. Choi et al.^23^ have found that the activation region binds to two histone acetyltransferases, p300 and CREB-binding protein (CREBBP also known as CBP). These interactions elevate the levels of acetylated histone H3 in nucleosomes around DUX4 binding sites, and through chromatin remodeling promote the expression of nearby genes. Recently, Vuoristo et al.^25^ showed that the activation region also binds to the Mediator complex through MED15, thereby suggesting an additional route by which the transcription machinery is recruited to the sites of DUX4-induced genes. A study by Ansseau et al.^26^ has identified and validated several interacting partners related to RNA processing including Complement C1q binding protein (C1qBP). They determined these interactions to be mediated by the N-terminal of DUX4, since they are also observed with shorter variants of the protein that lack the C-terminal. The interaction with C1qBP was also confirmed, by co-immunoprecipitation, at the endogenous level in FSHD muscle cells^15^, and recently by in situ proximity ligation assay on FSHD muscle sections^27^.

Despite these efforts, our knowledge of the DUX4 interactome is still partial. The unusual sequence features of DUX4 pose unique challenges to the most common approach to identify protein-protein interactions: Affinity Purification and Mass Spectrometry (AP-MS). Consequently, previous applications of AP-MS to DUX4^25,26^ have reported large sets of putative protein partners, from which it was hard to identify the few that are most functionally relevant. Therefore, key players in the early stages of DUX4 toxicity may have been missed. To rectify this situation, we present an AP-MS study of the DUX4 interactome, which we have specifically modified to account for the different aspects of DUX4 functions. Our results revalidate the interactions previously found, and reveal several new interaction partners that have direct relevance to DUX4 roles in human health and disease.

## Results

### DUX4 presents unique challenges for interactome analysis

The protein sequence of DUX4 presents a charge distribution that is among the most polarized to be found in any human protein. Under physiological conditions, the large number of arginine residues at the N-terminal and glutamic acid residues at the C-terminal lead to net charges of +17 and -15 (elementary charge units) at the respective terminal (Fig. 1A). The scan of the entire human proteome for the net charge in any stretch of 30 consecutive amino acids reveals that the two termini of DUX4 are located at the two extremes of the distribution (Fig. 1B). The unusual charge distribution poses unique challenges for the study of the DUX4 protein interactome by affinity purification and mass spectrometry (AP-MS). In particular, abundant proteins that bind non-specifically to DUX4 because of favorable electrostatic interactions may be erroneously classified as interacting partners.

In this study, we have taken several steps to mitigate these artifacts of the AP-MS measurements and provide a rigorous analysis of the DUX4 interactome. First, we employed different experimental controls tailored to the net charge of each terminal. Second, we used the high-affinity Strep-tag on the bait proteins. This allowed for high-salt washes that reduced the signal from non-specific interactions. Third, we placed the Strep-tag at the N-terminal of our bait proteins rather than at the C-terminal. As the very last residues of the DUX4 sequence were shown to be critical for the activation of its toxicity^23,24^, the location of the tag at the N-terminal minimizes interference to these functions. Fourth, we compared the protein abundances in the pulldown analyses to their endogenous cellular values in order to assess their enrichment in the pulldown, with higher enrichment values signifying a more likely interaction. Finally, we prolonged the expression times of DUX4 before the AP-MS analysis to 48 hours, in order to allow the cellular abundances of its induced proteins to reach detectable levels.

### Protein interactions associated with the C-terminal activation region of DUX4

In order to identify the protein interactions that are associated with the activation region of DUX4, we have performed AP-MS experiments of the full DUX4 protein against the negative control DUX4_ΔC construct (Table 1). To that end, we transfected HEK293 cells with plasmids for the constitutive expression of either DUX4 or DUX4_ΔC, and observed their increasing levels over a time course of 48 hours. Starting at 24 hours, the protein expression of several DUX4-induced genes, such as RFPL4B and ZSCAN4, was observed only in the DUX4-transfected cells. The cells were collected and lysed 48 hours post transfection, and the lysates were subjected to affinity purification by Streptactin resin. After the resin was washed, the proteins that remained bound to the resin were eluted and analyzed by mass spectrometry.

**Table 1.**
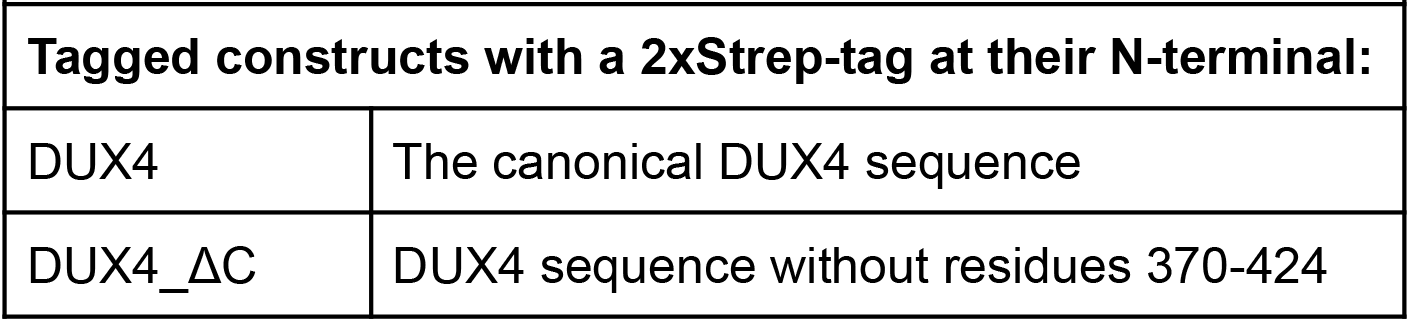
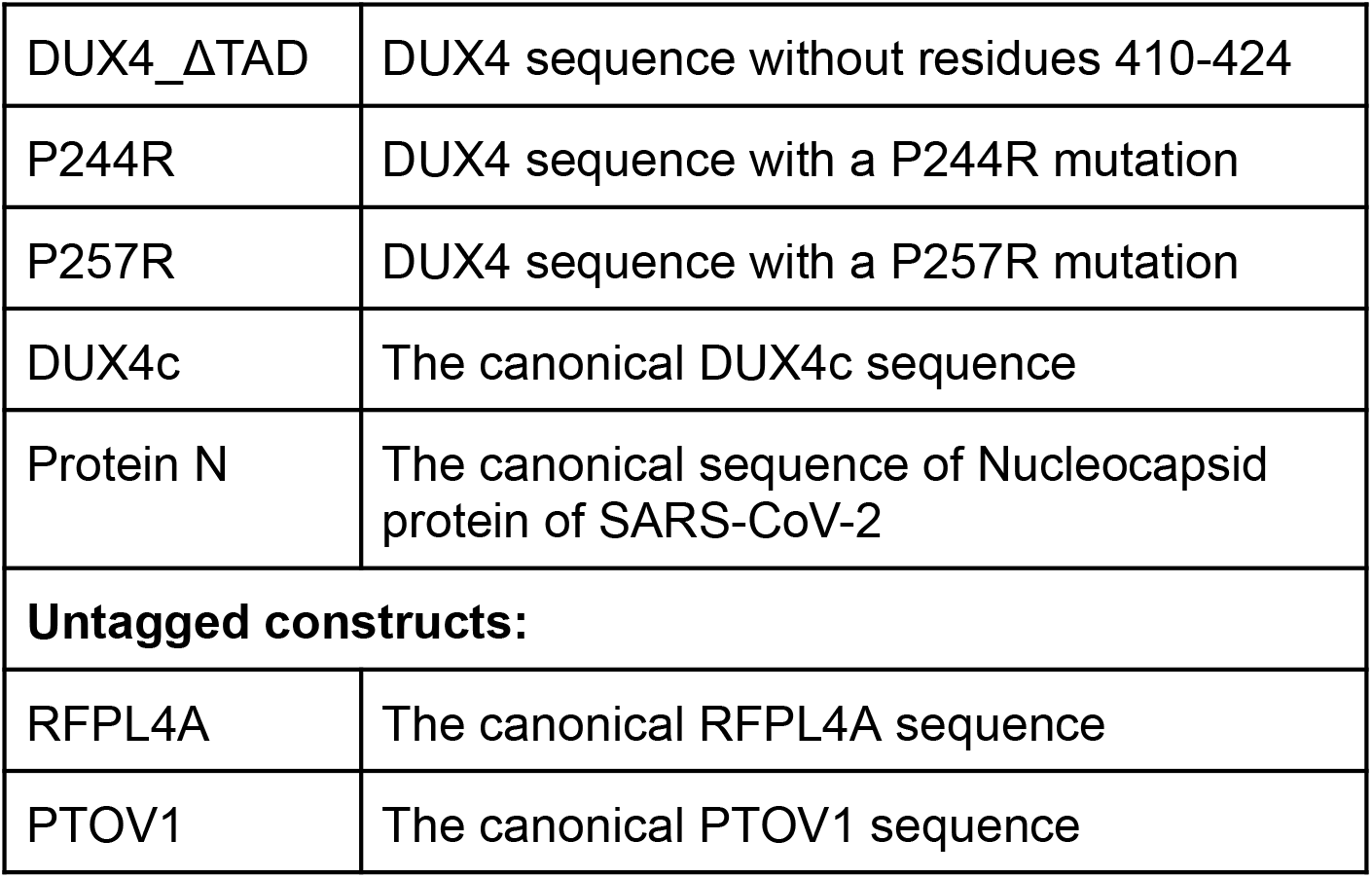
Protein constructs used in this study.

| Table 1. Protein constructs used in this study. |  |
| --- | --- |
| Tagged constructs with a 2xStrep-tag at their N-terminal: |  |
| DUX4 | The canonical DUX4 sequence |
| DUX4_ΔC | DUX4 sequence without residues 370-424 |
| DUX4_ΔTAD | DUX4 sequence without residues 410-424 |
| P244R | DUX4 sequence with a P244R mutation |
| P257R | DUX4 sequence with a P257R mutation |
| DUX4c | The canonical DUX4c sequence |
| Protein N | The canonical sequence of Nucleocapsid protein of SARS-CoV-2 |
| **Untagged constructs:** |  |
| RFPL4A | The canonical RFPL4A sequence |
| PTOV1 | The canonical PTOV1 sequence |

The AP-MS analysis (Fig. 2, Table S1) revalidates the known protein interactions associated with DUX4 activation. Both p300 and CREBBP were highly enriched only in the DUX4 pulldown. Because these proteins are endogenously expressed, and are not DUX4-induced, their significant enrichment reaffirmed their direct and strong binding affinity with the activation region^23^. The H3.X and H3.Y histone variants were also significantly enriched in the DUX4 pulldown. Resnick et al.^28^ have shown that these variants are incorporated into the chromatin of transcribing genes that are induced by DUX4. In accord with that study, peptides that are unique to H3.X and H3.Y were readily detectable only in the DUX4-transfected cells, both at the whole-cell lysate and the affinity-enriched pulldown. We note however that all these peptides mapped to sequence regions that are identical between the two variants. Hence, we cannot determine what is the abundance ratio between H3.X and H3.Y, and they are collectively referred to as H3.X/Y. The most plausible explanation for the enrichment of these histone variants would be an indirect interaction of the DNA-binding domains of DUX4 with DNA in chromatin that is enriched with H3.X/Y.

**Figure 2.**
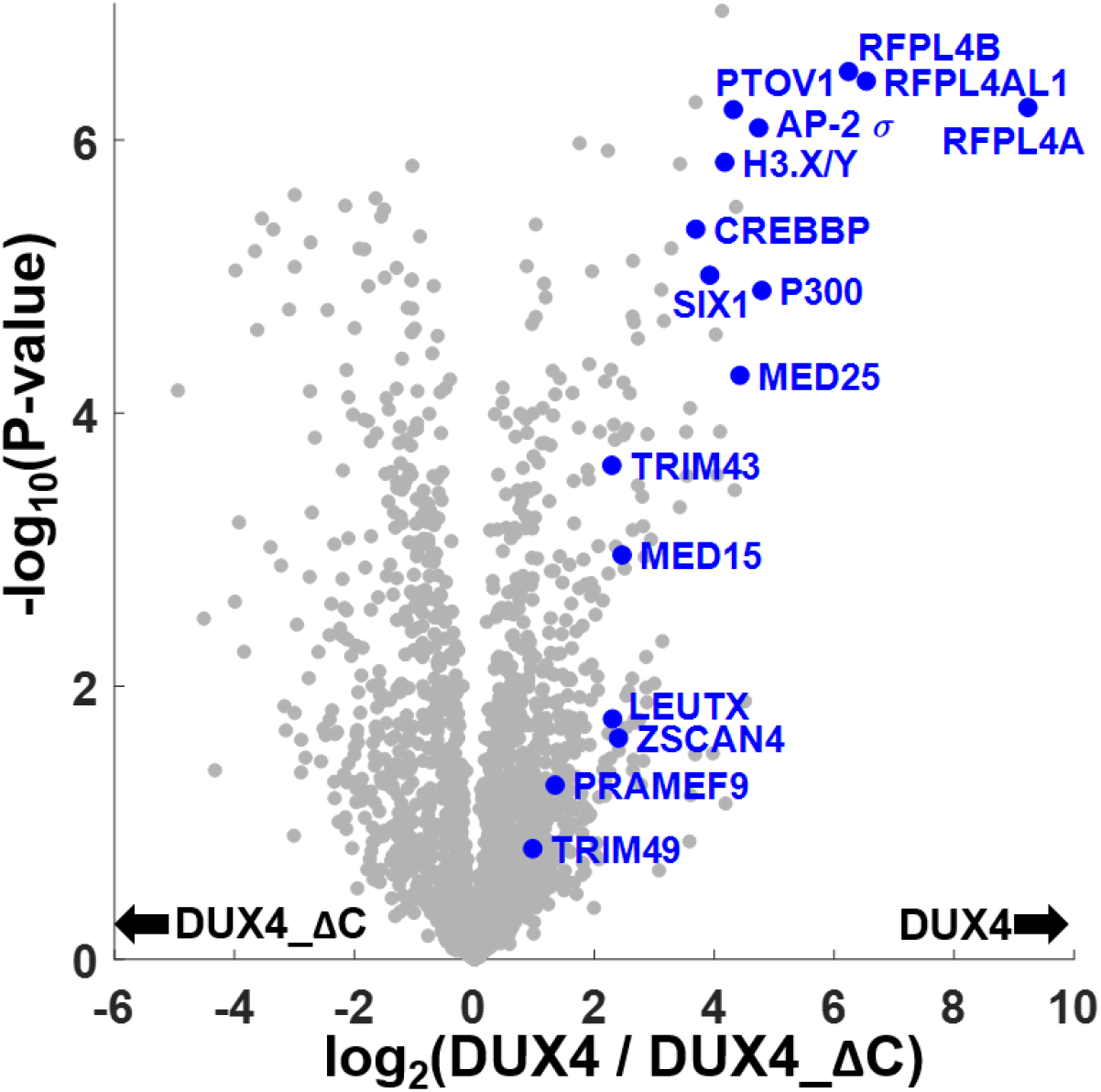
Protein interactions associated with the C-terminal activation region of DUX4. The affinity purification and mass spectrometry (AP-MS) results from HEK293 cells that express either tagged DUX4 or DUX4_ΔC. Affinity purification was performed 48 hours following transfection with the expression plasmids. The x-axis measures the abundance ratio of a specific protein in the pulldowns of DUX4 relative to DUX4_ΔC. The y-axis measures the confidence level assigned to the abundance ratio. The right side of the plot features proteins that are more abundant in DUX4 pulldowns, with certain proteins-of-interest labeled in blue. Note the significant presence of DUX4-induced genes, such as RFPL4 and LEUTX, in the DUX4 pulldowns. The affinity purification experiments were performed in biological quadruplicates.

We were surprised to see the presence of multiple DUX4-induced proteins in the DUX4 pulldowns. The most prominent signal belonged to members of the Ret finger protein-like 4 family (RFPL4A, RFPL4AL1, and RFPL4B), but TRIM43, TRIM49, LEUTX, PRAMEF9, and ZSCAN4 were also readily detectable. The fact that these proteins are not endogenously expressed in HEK293 cells, raises the concern that their appearance at the right side of the plot may be an artifact of the DUX4 induction (and its lack thereof in DUX4_ΔC cells). For example, non-specific binding of the DUX4-induced proteins to the Strep-Tactin resin will lead to similar AP-MS results. To address this concern, we looked in more detail at the enrichment of these proteins by the purification process (Table 2). To that end, we performed standard proteomics analyses of the whole-cell lysates prior to the Strep purification. We then sorted the detected proteins from each sample according to their abundance (i.e., the most abundant protein had a rank of 1). The ranking of the proteins at the lysate level can now be compared with their ranking following the Strep purification. The largest increase in rank belongs to RFPL4A and RFPL4AL1, which are not detectable at the lysate level, but become highly abundant (ranks of 21 and 73, respectively) after the Strep purification. Approximately 3,500 different proteins are detected in a lysate sample, implying that the rank of undetected proteins is lower than 3,500 in the whole-cell lysate. Hence, such a large increase in the rank (i.e. from >3,500 to 21) is most likely the result of a specific interaction between DUX4 and RFPL4A, rather than the outcome of a non-specific binding between RFPL4A and Strep-Tactin resin. Of note are large increases in rank also for RFPL4B, LEUTX, and PRAMEF9, again arguing for a specific interaction between each of these proteins and DUX4. On the other hand, TRIM43, TRIM49, and ZSCAN4 show only moderate improvement in rank, and therefore a specific interaction between them and DUX4 is less certain and will not be further discussed.

**Table 2.** Abundance ranks of selected proteins 48 hours after the transfection with either DUX4 or DUX4_ΔC plasmids. The most abundant protein in a certain sample has a rank of 1. The results are the averages from quadruplicates for both the Strep-affinity purifications and the whole-cell lysates. n.d. – The protein was not detected in the mass spectrometry analysis, which typically identifies 3,500 different proteins.

| Gene name | UNIPROT accession code | Abundance rank after Strep purification |  | Abundance rank at the cellular lysate level |  |
| --- | --- | --- | --- | --- | --- |
|  |  | DUX4 | DUX4_ΔC | DUX4 | DUX4_ΔC |
| Interactions associated with the activation region: |  |  |  |  |  |
| RFPL4A | A6NLU0 | 21 | n.d. | n.d. | n.d. |
| RFPL4AL1 | F8VTS6 | 73 | n.d. | n.d. | n.d. |
| RFPL4B | Q6ZWI9 | 113 | n.d. | 1496 | n.d. |
| P300 | Q09472 | 323 | n.d. | 3644 | 3176 |
| CREBBP | Q92793 | 666 | n.d. | n.d. | n.d. |
| PTOV1 | Q86YD1 | 305 | 2377 | n.d. | n.d. |
| MED25 | Q71SY5 | 452 | 2633 | 3686 | 3613 |
| H3.X/Y | P0DPK5;<br>P0DPK2 | 303 | n.d. | 1899 | n.d. |
| SIX1 | Q15475 | 571 | 1109 | n.d. | n.d. |
| AP-2-σ | P53680 | 396 | n.d. | 2173 | 1803 |
| AP-2-μ | Q96CW1 | 168 | 1359 | 1013 | 1121 |
| MED15 | Q96RN5 | 835 | 1646 | n.d. | n.d. |
| TRIM43 | Q96BQ3 | 1360 | n.d. | 2061 | n.d. |
| LEUTX | A8MZ59 | 1137 | n.d. | n.d. | n.d. |
| PRAMEF9 | Q5VWM5 | 1096 | n.d. | n.d. | n.d. |
| TRIM49 | P0CI25 | 1491 | n.d. | 2030 | n.d. |
| ZSCAN4 | Q8NAM6 | 1070 | n.d. | 1912 | n.d. |
| N-terminal interactions: |  |  |  |  |  |
| C1QBP | Q07021 | 2 | 2 | 98 | 73 |
| DNAJC9 | Q8WXX5 | 7 | 6 | n.d. | n.d. |
| PARP-1 | P09874 | 84 | 42 | 101 | 102 |
| XRCC5 | P13010 | 40 | 27 | 134 | 121 |
| XRCC6 | P12956 | 89 | 55 | 229 | 194 |
| H2AFX | P16104 | 120 | 57 | 553 | 621 |
| PRMT5 | O14744 | 174 | 62 | 697 | 647 |
| WDR77 | Q9BQA1 | 74 | 24 | 797 | 849 |
| Tagged proteins: |  |  |  |  |  |
| DUX4 | Q9UBX2 | 1 | -- | 173 | -- |
| DUX_ΔC | -- | -- | 1 | -- | 51 |
| Background proteins: |  |  |  |  |  |
| Actin | P60709 | 23 | 29 | 1 | 1 |
| EF1A1 | P68104 | 34 | 69 | 2 | 2 |
| HSP90B | P08238 | 80 | 88 | 3 | 3 |

Two transcription activators, which were previously not reported as interactors of DUX4, are prominent in the AP-MS results: Prostate Tumor-Overexpressed gene 1 (PTOV1) and Mediator of RNA polymerase II transcription subunit 25 (MED25). Examination of their rank change between the whole-cell lysate and the pulldown (Table 2) suggests a direct interaction with DUX4, with an enrichment factor that is on par with p300. This interaction is likely mediated by a common homologous region, the PTOV-domain^29^, which is present in both proteins. While the cellular functions of PTOV1 are not well-characterized, MED25 is an established component of the Mediator complex. Hence, the interaction indicates a direct mechanism by which DUX4 can recruit the Mediator complex, and subsequently RNA polymerase II, to promote the transcription of induced genes.

Additional proteins in the AP-MS results may have relevance to DUX4 toxicity. First, Homeobox protein SIX1 (SIX1) plays a role in myogenesis, including adult muscle regeneration, and in the modulation of the fast-twitch glycolytic phenotype^30^. SIX1 was also reported to be essential for myoblast differentiation and fusion^31^. Second, the entire AP-2 (adaptor protein 2) complex is significantly enriched in the DUX4 pulldown, with subunits sigma, mu, gamma, alpha-1 and 2 having the most prominent signals. AP-2, acting in clathrin-dependent endocytosis, may be relevant to DUX4 toxicity because of its involvement in muscle cell fusion^32^. The molecular basis for this interaction may be the DUX4 KIX-binding domain in the activation region^33^, which corresponds to the endocytosis signal motif [ED]-X-X-X-L-[LI] that is recognized by AP-2 sigma^34^. We recently detected DUX4 both in nuclei and cytoplasm of myogenic progenitors in FSHD muscle sections^27^, thereby suggesting that cytoplasmic DUX4 may also directly interact with AP-2 in those cells.

### Protein interactions associated with the N-terminal DNA-binding region of DUX4

In order to identify protein partners that bind exclusively outside the activation region of DUX4, we have performed AP-MS analysis of DUX4_ΔC against a control of cells transfected with an empty plasmid (Figure 3A, Table S4). Because of the large net positive charge of the N-terminal, we were concerned that some of the enriched proteins, such as the negatively-charged tubulin TUBB8 protein, may be the result of nonspecific electrostatic interaction. To account for such possible artifacts, we performed a second AP-MS analysis in which the control arm was a different protein of significant positive surface charges (Figure 3B). We chose the nucleocapsid protein (Protein N) of SARS-CoV-2 that is positively charged, but is otherwise unrelated to any aspect of the DUX4 toxicity^35^. The new control reduced the enrichment signal of several previous hits, resulting in a more specific set of N-terminal protein partners. Similar analyses with the full-length DUX4 pulldowns gave essentially the same set of N-terminal interactions (Figure S1).

**Figure 3.**
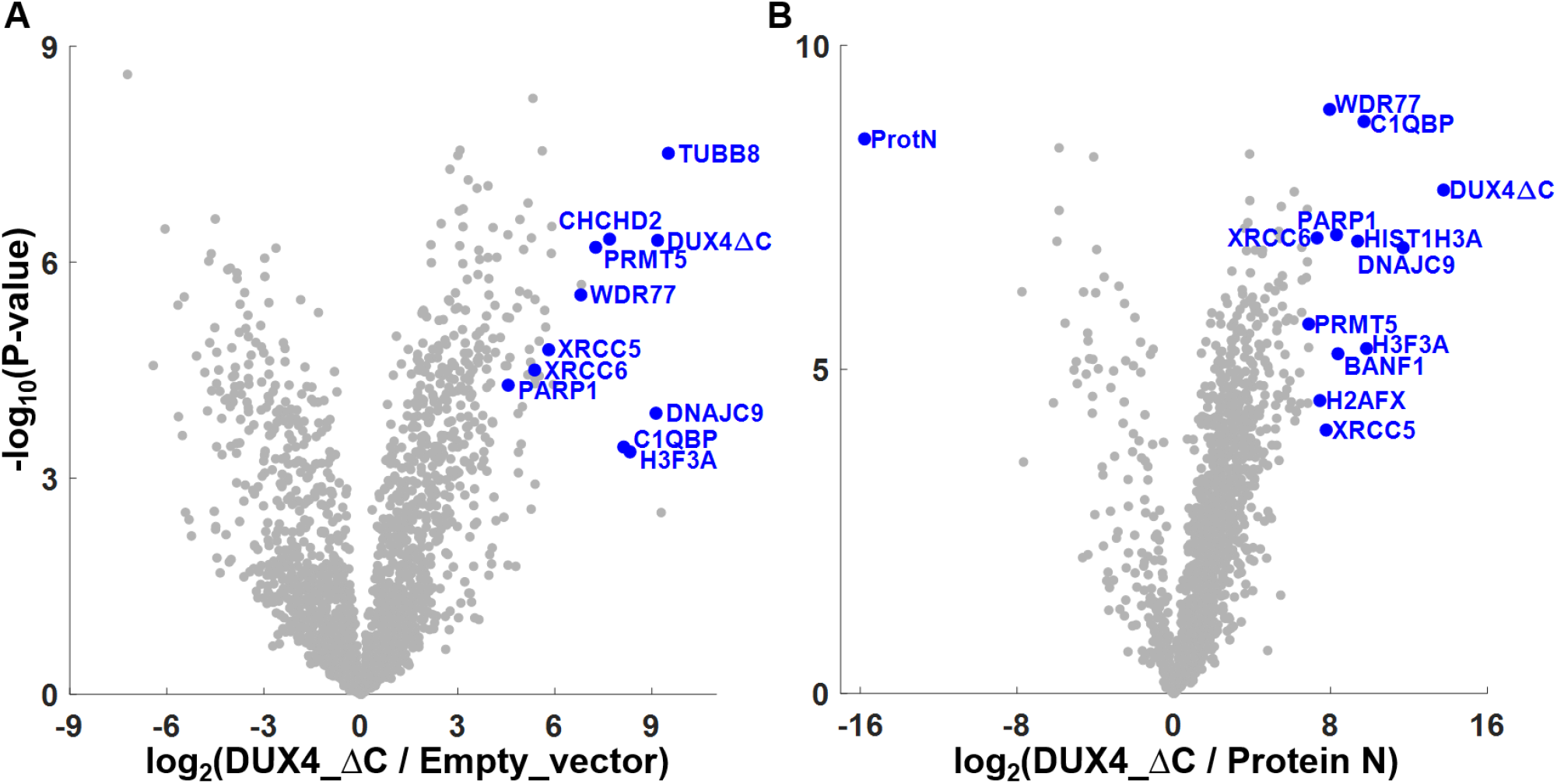
Protein interactions associated with the N-terminal DNA-binding region of DUX4_ΔC. Results of AP-MS analysis from HEK293 cells that express either DUX4_ΔC, an empty transfection vector, or the Nucleocapsid protein of SARS-CoV-2 (Protein N). The affinity purification experiments were performed in biological quadruplicates.

One of the strongest interactions of the N-terminal is with the Complement component 1 Q subcomponent-binding protein (C1qBP), a highly acidic protein with multiple cellular localizations and functions^36^. Indeed, the DUX4-C1qBP interaction was previously reported *in-vitro*^15,26^ and *in-situ* in human muscle sections^27^. In non-human myoblasts, C1qBP loss-of-function inhibits their proliferation and differentiation, leading to their death^37^. Hence, the strong interaction with DUX4 may interfere with normal C1qBP functions in a similar manner. Another strong interaction is with DnaJ homolog subfamily C member 9 (DNAJC9), which to the best of our knowledge was not previously reported for DUX4. DNAJC9 is an established histone chaperone involved in histone deposition during DNA replication and transcription^38^. Hence, this interaction may facilitate the deposition of H3.X and H3.Y histone variants around the sites of DUX4 binding. Of note is the high enrichment of DNAJC9 in the DUX4 pulldown compared to its general cellular abundance (Table 2), which suggests a direct binding with DUX4 rather than an indirect interaction that is mediated via the DNA binding.

We observe that the N-terminal interactions are highly enriched with proteins that are involved in the regulation of DNA repair following double-strand breaks (DSBs). These include well-known DSB sensors such as the XRCC5/6 complex (also known as the Ku complex), PARP1, and H2AFX: the H2AX histone variant acting as a DSB sensor when phosphorylated at a specific site^39,40^. However, other top interactions are implicated in DSB pathways as well. BANF1 was reported to interact with PARP1 leading to defective DNA repair^41^. C1qBP was reported to regulate the MRN complex that plays critical roles in DSB repair response^42^. PRMT5 is involved in various biological processes either with WDR77, forming part of the methylosome complex modifying several histones and spliceosomal proteins, or with other partners to up-regulate genes involved in DNA repair^43,44^. PRMT5 was also found to be required in skeletal muscle for MyoD-dependent differentiation of muscle stem cells and their maintenance in adult regeneration^45,46^.

### Domain-level localization of the DUX4 interactions with RFPL4A and PTOV1

In order to better characterize the interactions of DUX4 with RFPL4A and PTOV1, we sought an experimental setup that decouples their protein expression from that of DUX4. To that end, we overexpressed in HEK293 cells either RFPL4A or PTOV1, lysed the cells, and saved the lysates as an external source for these two proteins in an untagged and an unpurified form. In parallel, we expressed as before the Strep-tagged DUX4 and DUX4c constructs, lysed the cells, and spiked the DUX4/4c lysates with a small volume (1/100 v/v) of either the RFPL4A or PTOV1 lysates. We then performed affinity purification and mass spectrometry analyses of the spiked lysates as described above. The control arms for the AP-MS experiments were lysates from mock-transfected cells that were similarly spiked with the RFPL4A or PTOV1 lysates (Figure 4, Table S5). Note that the DUX4c construct is used here for a similar purpose as DUX4_ΔC. The sequences of DUX4c and DUX4 are identical in their first 342 residues, and DUX4c lacks the activation region of DUX4.

**Figure 4.**
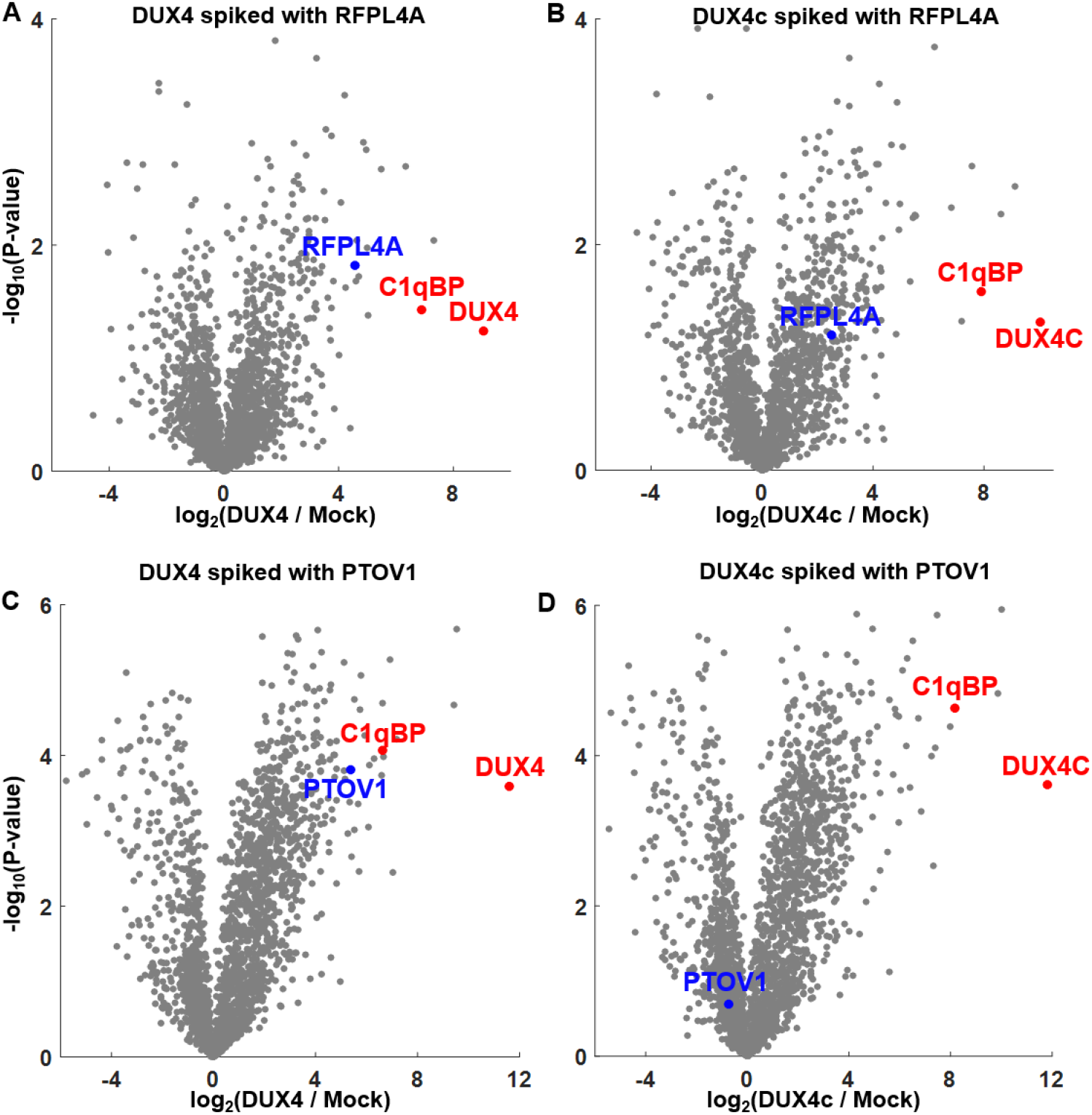
AP-MS analyses of spiked lysates. Purifications of Strep-tagged DUX4 or DUX4c from lysates spiked with either RFPL4A **(A,B)** or PTOV1 **(C,D)**. RFPL4A is being pulled down by both DUX4 and DUX4c, indicating an interaction with their identical N-terminal sequence. PTOV1 is only being pulled down by DUX4, indicating a direct interaction with the C-terminal activation region. The location of C1qBP, an established N-terminal interactor, is shown for reference. The affinity purification experiments were performed in biological duplicates.

The results show that RFPL4A is being pulled down equally by both DUX4 and DUX4c (Figure 4A,B). In fact, the apparent two-fold higher intensity of RFPL4A in the DUX4 pulldown compared to that of DUX4c stems from the additional DUX4-induced expression of RFPL4A, which is absent in the DUX4c cells. Hence, this pulldown pattern localizes the interaction to the first 342 residues of the DUX4 sequence that are identical in DUX4c. In contrast, PTOV1 is only being pulled down by DUX4 and not DUX4c (Figure 4C,D). This pulldown pattern indicates a direct interaction between PTOV1 and the C-terminal activation region of DUX4.

### All-atom modeling of the DUX4-RFPL4A interaction

For structural characterization of the DUX4-RFPL4A interaction we analyzed it by cross-linking and mass spectrometry (CL-MS)^47^. To that end, we formed the DUX4-RFPL4A complex by mixing lysates from cells overexpressing the two respective proteins. We cross-linked the complex in turn with several cross-linking reagents (BS^3^, Formaldehyde, and EDC) that target different amino acid side chains. We then purified the cross-linked complex via the strep tag on DUX4, analyzed it by mass spectrometry, and performed searches for the respective cross-linking chemistries (see Methods). The two cross-links that were identified (Figure 5A) are: BS^3^ cross-link between DUX4-Y_241_ and RFPL4A-K_177_, and formaldehyde cross-link between DUX4-R_258_ and RFPL4A-K_177_. Both cross-links had excellent MS/MS fragmentation evidence to support their identification (Figure S2). Because the two cross-links relied on different chemistries, their convergence to the same regions on both proteins supports the relevance of these regions for the interaction.

**Figure 5.**
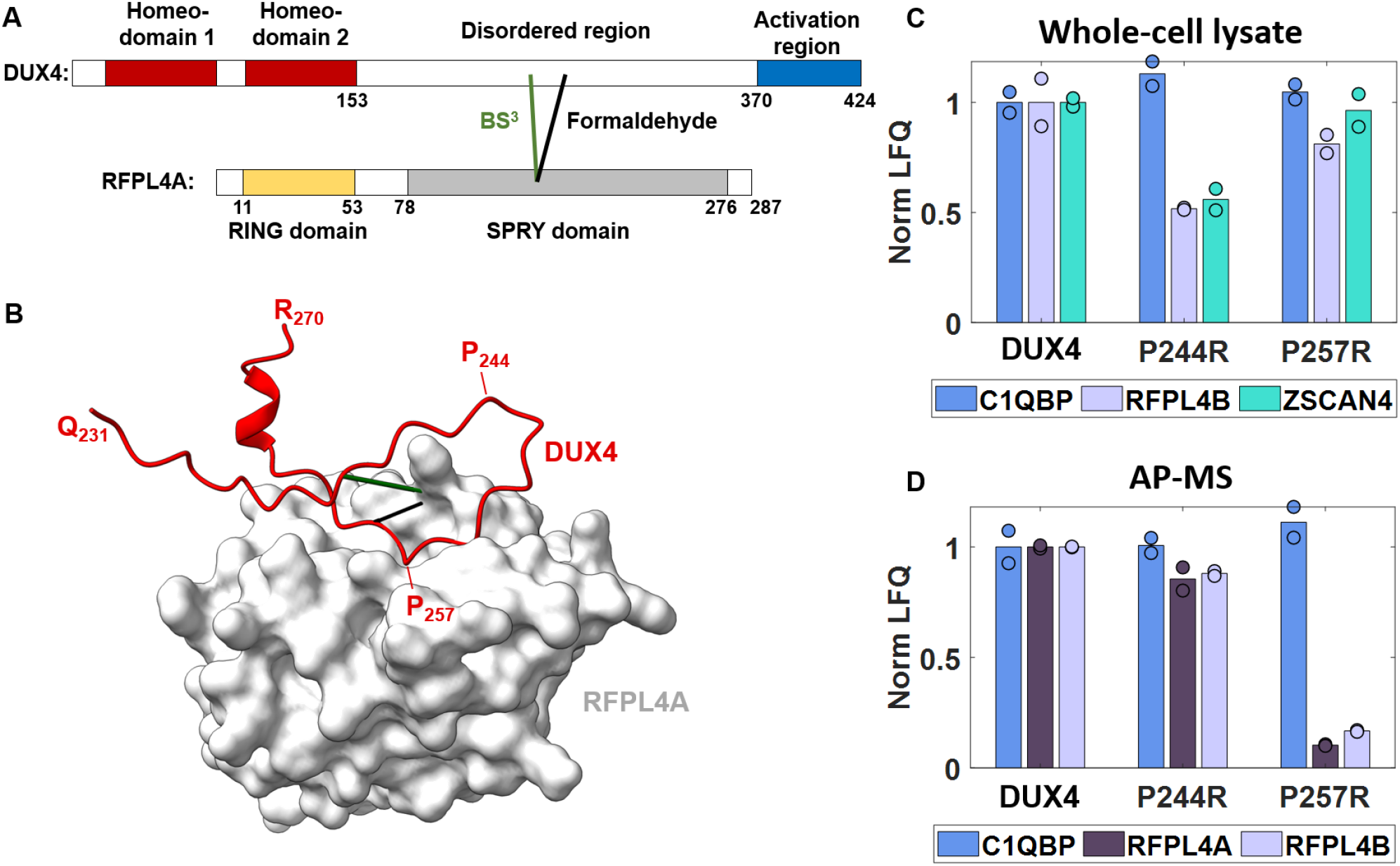
All-atom modeling and validation of the DUX4-RFPL4A interaction. **(A)** Cross-linking and mass spectrometry analysis of the interaction by two different reagents (BS^3^ and Formaldehyde) identified two cross-links (solid lines) that are mapped onto the protein sequences. **(B)** An all-atom computational model by AlphaFold-Multimer of the SPRY domain of RFPL4A (gray) interacting with the DUX4 disordered region surrounding the cross-links (red). The model agrees with the cross-links and places Pro_257_ of DUX4 at the center of the interaction, whereas nearby Pro_244_ is not involved. **(C,D)** Validation of the model by deleterious point mutations. The DUX4, P244R, and P257R constructs activate in cells the expected genes (RFPL4B and ZSCAN4) in similar quantities (Whole-cell lysate), yet the P257R protein has lost its ability to pull down RFPL4 proteins by Strep purification (AP-MS). The DUX4-C1qBP interaction that does not involve the disordered region serves as a positive control. All values are normalized compared to the corresponding average values of the DUX4 construct. Experiments were performed in biological duplicates.

Next, we employed AlphaFold-Multimer (AFM)^48^ to obtain an all-atom model of the interaction. The input sequences for AFM were the full-length sequence of RFPL4A and residues 231-270 of DUX4. The sequence of DUX4 was trimmed to assist the convergence of AFM by reducing the search space, and was chosen to encompass the identified cross-linked residues. Of note, apart from their role in trimming the DUX4 input sequence, the cross-links were not otherwise used by AFM. The best scoring model had good confidence scores (Figure S3), and docked residues 253-260 of DUX4 into the substrate binding groove of the SPRY domain of RFPL4A, in full agreement with the two cross-links (Figure 5B, Supplementary Dataset DS1). According to the model, the interaction is centered around a pair of DUX4 residues, Ala_256_Pro_257_, that forms favorable hydrophobic interactions deep within the binding groove. Additional support for the correctness of the model can be found in the study of Grütter et al.^49^, which solved the crystal structure of RFPL4A with a substrate peptide bound to the SPRY domain. Also in the crystal structure, an Ala-Pro motif is central to the interaction and binds in a similar manner.

The Ala-Pro motif occurs four times in region 231-270 of DUX4. Therefore, further validation is required that the correct Ala-Pro motif was indeed indicated by AFM. To that end, we tested two mutated DUX4 constructs (Table 1): (i) the P257R that targets the Ala-Pro motif predicted at the center of the interaction, and (ii) the P244R that targets a nearby motif predicted to be uninvolved (Figure 5B). Both mutation constructs resulted in active DUX4 that led to the detection of the expected induced genes by mass spectrometry analyses of the whole-cell lysates (Figure 5C, Table S6). However, the ability of the P257R construct to pull down members of the RFPL4 family by Strep-tag purification is nearly abolished, whereas the P244R construct behaves similarly to the canonical DUX4 sequence (Figure 5D, Table S6). We conclude that the all-atom model generated by AFM indeed describes the structural basis for the interactions of DUX4 with members of the RFPL4 protein family.

### All-atom modeling of the DUX4-PTOV1 interaction

A similar experimental/computational pipeline was conducted to model and validate the DUX4-PTOV1 interaction. We formed the interaction complex by mixing lysates from cells overexpressing the two respective proteins, and performed CL-MS. Because the previous data indicate that PTOV1 interacts with the DUX4 activation region (Figure 4), adjustments were made in the CL-MS analysis. Specifically, the lack of lysine and arginine residues in the activation region excludes the use of BS^3^ and formaldehyde reagents for cross-linking, and prevents the digestion of this region by trypsin. These impediments were rectified by employing EDC cross-linking reagent that targets both lysine and acidic side chains, and by digesting the proteins with a combination of trypsin and GluC proteases. We identified 25 relevant EDC cross-links, nearly all of them between the second half of the activation region of DUX4 (residues 390-424) and both PTOV1/MED25 domains in the PTOV1 protein (Figure 6A, Table S7). The fact that both PTOV1/MED25 domains can independently bind the activation region may be a contributing factor for the particular high affinity of the interaction.

**Figure 6.**
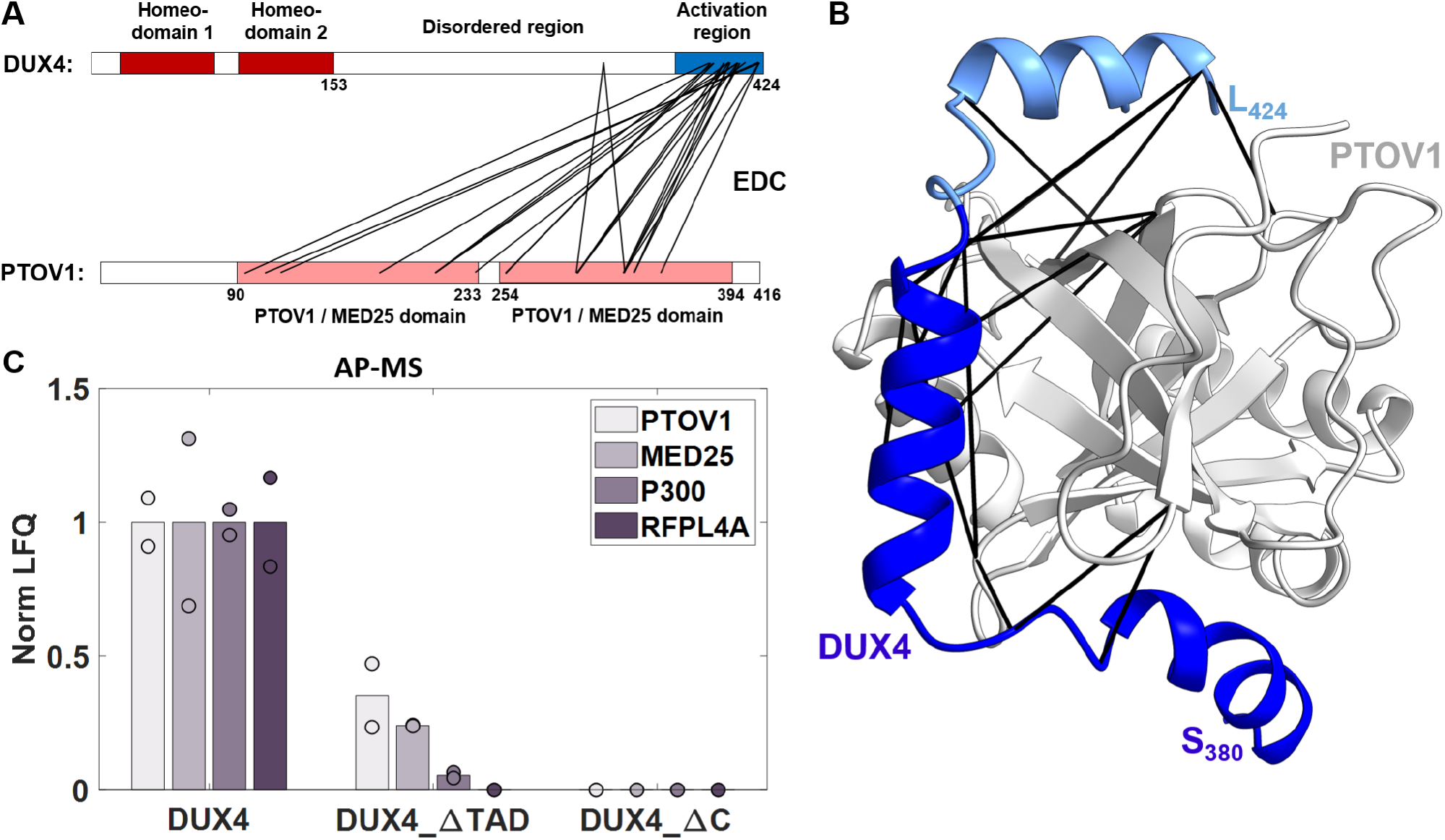
All-atom modeling and validation of the DUX4-PTOV1 interaction. **(A)** Cross-linking and mass spectrometry identified 25 EDC cross-links (black lines) between the second half of the activation region of DUX4 and the globular PTOV1 domains. **(B)** The AlphaFold-Multimer model wraps the DUX4 activation region (blue) along a groove on the surface of the PTOV1 domain (gray) in a manner that is consistent with the cross-link set (black). Note the extensive interface area of the interaction (∼2,800 Å^2^). A region of DUX4 that was deleted in a follow-up validation experiment (DUX4_ΔTAD) is marked in light blue. **(C)** AP-MS values of four proteins from cells expressing three different constructs (Table 1). DUX4_ΔTAD retains some of its ability to bind PTOV1 and MED25, in accordance with the model. All values are normalized compared to the average DUX4 value. Experiments were performed in biological duplicates.

We employed the cross-link information to trim the input sequences of AFM to residues 371-424 of DUX4 (the activation region) and residues 245-416 of PTOV1 (the second PTOV1 domain). The resulting models converged to an extended interface of the DUX4 activation region wrapped along a groove on the surface of the second PTOV1 domain (Figure 6B, Supplementary Dataset DS2). By employing the high sequence homology between the two PTOV1 domains, it is straightforward to map all the cross-links involving the first domain onto their corresponding residues in the second domain. Mapping this combined cross-link set onto the AFM model reveals a remarkably good agreement. Although the model predicted an interaction interface between the PTOV1 domain and the entire activation region (residues 371 to 424), the cross-links coverage is narrower (residues 390 to 424). The latter sequence region is therefore more likely to be representative of the true extent of the interaction. Even this more restricted interface is still extensive, burying an exposed surface area of 2,800 Å^2^ and wrapping around more than half the circumference of the PTOV1 domain.

We further validated the AFM model by testing the binding of PTOV1 to a truncated DUX4 construct that lacks the last 15 residues of the activation region (DUX4_ΔTAD, Table 1). To that end, we compared the abundances of several proteins-of-interest in the AP-MS results from DUX4_ΔTAD, DUX4_ΔC, and DUX4 experiments (Figure 6C, Table S8). The truncation substantially reduced, but did not eliminate, the ability of the DUX4_ΔTAD construct to bind to PTOV1. Such reduced affinity is in perfect accord with our structural model in which the TAD truncation comprises only half of the binding interface. Interestingly, the truncation also reduced, but did not eliminate, the binding of p300. This strongly suggests that the binding of p300, PTOV1, and MED25 onto DUX4 largely overlap along its sequence. We also observed that the TAD truncation inactivated DUX4 in the context of HEK293 cells so that downstream genes were not induced (such as RFPL4’s). Overall, these results support the structural model, which in turn provides the structural basis for the critical role of the last 15 residues of DUX4 to its toxicity as previously reported^24^.

## Discussion

This study employed an AP-MS methodology that was specifically tailored to the unique features and functions of DUX4. The approach successfully delineated a narrow set of proteins that showed particularly strong binding to DUX4. This set included all the previously verified interactions (CBP/p300, C1qBP, and the enrichment of H3.X/Y in the chromatin), as well as new identifications (mainly RFPL4s, PTOV1, and MED25) that paint a more complex picture of DUX4 activation and regulation. In particular, MED25 is a hub protein that, in addition to its established role in the Mediator complex, also interacts with the CBP/p300 acetyltransferases and the retinoic acid receptor^29^. Indeed, the ability of DUX4 to directly recruit the Mediator to the sites of induced genes further explains its potency as a transcription factor.

The interaction with MED25 is in accord with Vuoristo et al.^33^ that have also found evidence for the binding of DUX4 to the Mediator complex. However, the AP-MS signal indicates that the binding to Mediator is facilitated through MED25 and not the adjacent MED15 subunit that was previously suggested^25^. Interestingly, MED25 and p300 appear to bind to a largely overlapping stretch of the DUX4 activation region. This is evident by our deletion study of the last fifteen residues of DUX4 that similarly impaired the binding of both proteins (Figure 6C). The overlap suggests a versatile and serial role of DUX4 in activation (Figure 7). First, it acts as a pioneering factor that opens the chromatin through the binding of p300^23^. Next, p300 dissociates and is replaced by MED25 on the same activation region to recruit the Mediator and subsequently RNA Polymerase II.

**Figure 7.**
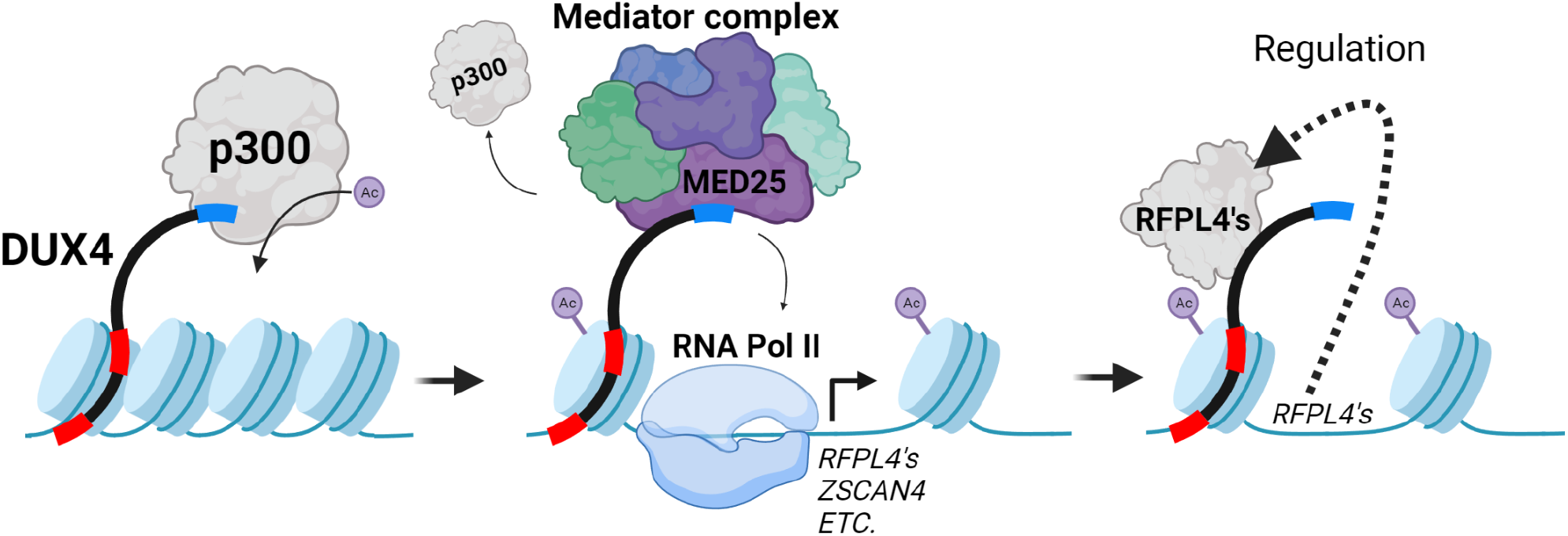
An extended model of DUX4 activation.

DUX4 interacts strongly with all members of the RFPL4 protein family, which are also among its most prominent induced gene targets. Three lines of evidence verified that these interactions are not an artifact. First, the high enrichment factor of RFPL4s in DUX4 pulldowns compared to their low abundance in the whole cell lysate is only consistent with a specific interaction (Table 2). Second, DUX4c (that is unable to induce RFPL4 expression) is also able to pull down RFPL4A that was externally added to the lysate (Figure 4B). Finally, a combination of CLMS and computational modeling have localized the interaction to the disordered region of DUX4 around Pro_257_, which was shown by point mutation to be critical for the interaction (Figure 5C). The strong DUX4-RFPL4 interaction forms a feedback loop with probable implications for DUX4 function. RFPL4 is an E3 ubiquitin ligase that works with the E2 ubiquitin-conjugating enzyme UBE2A^50^. Therefore, we hypothesize that this is a negative feedback loop that eventually inhibits DUX4 activity by ubiquitination and subsequent degradation (Figure 7). Such a mechanism may underlie the pulse-like expression of DUX4 that is observed both during embryogenesis^51^ and in myotubes from patients with FSHD^6^. We believe that this merits further research since the feedback loop may explain the large variance in the FSHD severity phenotype that is observed between individuals that otherwise have a similar DUX4 genotype^52^.

The large extent of the interaction between the activation region and the PTOV1/MED25 domain was unexpected. Analysis of evolutionary conserved residues between human DUX4 and its mouse orthologue shows that only residues 410 to 424 are conserved between the species^53^. These fifteen residues also harbor motifs found in other trans-activation domains (TADs). Yet, the cross-links and structural model report on a more extensive interaction that spans residues 390 to 424. This is further verified by the DUX4_ΔTAD construct that retains partial capability to bind to the PTOV1/MED25 domain (Figure 6C), probably through the remaining residues 390 to 409. We find the extended interacting region to be fortuitous from the viewpoint of future therapeutic strategies for FSHD. First, it presents more potential targets for inhibitory molecules that could interfere with the interaction. More importantly, it involves DUX4-unique regions preceding the TAD motif that may be therapeutically targeted without affecting the other TAD motif interactions in the cell.

The AP-MS analysis found that the group of interacting partners with the N-terminal of DUX4 is highly enriched with proteins involved in the DNA damage response. This is intriguing, since multiple studies have reported that DUX4 induces DNA damage^12,15,54^, but to our knowledge none refers to inhibition of the DNA repair pathway or the DNA damage response. We believe that further investigation into these interactions is merited as they may impact the normal functions of key elements of DNA repair, thereby bringing about a new toxicity pathway that operates at the protein level.

On a technical note, we think that the structural pipeline employed here demonstrates an important synergy between CLMS and deep-learning approaches. The information from the cross-links was used twice in the process. First, to shorten the sequence inputs to the computation so that the likelihood for a correct and converged ensemble of models is improved. Then, by selecting the best-fitting model from the ensemble. The computational modeling, in turn, provides atomistic details that cannot be achieved by the lower resolution inherent to CL-MS. The resulting all-atom model is then well-suited to design follow-up verification experiments. We foresee that this approach could be particularly useful to study other interactions that involve largely disordered regions such as those inherent to DUX4.

## Methods

### Plasmids

The expression vector for the canonical DUX4 and DUX4c, fused to Strep-tags at their N-terminal, were generated on the basis of the pFN21A-DUX4 and -DUX4c vectors we previously used^26^. These vectors were linearized using the NheI and PvuI enzymes to remove the existing HaloTag. The fragments were separated by gel electrophoresis and the ∼5-kb band purified with a DNA gel extraction kit (Machery-Nagel). The In-Fusion Snap Assembly (Takara Bio) was used for cloning. The insert corresponds to the oligonucleotide sequence 5’-TCACTATAGGGCTA*GCcgccAccATGG*CGGGG<u>tctgcttggtcacacccacagttcgagaag</u>ggaggt ggatcaggcggaggcagcggaggc<u>tctgcttggtcacacccacagttcgagaag</u>ggaggatcaggatcaggcGCGAT CGCCATGGCCCTCCC-3’ (in double-strand with PAGE purification, Integrated DNA Technologies). The insert contains two Strep tags (underlined) separated by a flexible linker sequence. Another flexible linker was also added between the second Strep tag and the DUX4/4c coding sequences, as well as the human consensus Kozak at 5’ (in italic) to optimize translation of the fusion protein. The sequences at each end (uppercase) correspond to the homology sequences required for assembly. Before the cloning reaction, the insert was pre-heated 5 min at 80°C and maintained 3 min on ice and a 1:10 insert:vector molar ratio was used for assembly following manufacturer instructions. The production of Strep-DUX4 and -DUX4c proteins were validated by Western blotting on extracts of cells transfected with each construct.

The other plasmids described in Table 1 were generated by insertion of point mutations or stop codons using the high fidelity Phusion polymerase (product #M0530L; New England Biolabs), and verified by sequencing. The expression plasmid of the Nucleocapsid protein of SARS-CoV-2 (Protein N) was previously described^35^. Expression plasmids in human cells of untagged RFPL4 and PTOV1 were purchased from Sino Biological (products HG25044-UT and #HG20551-UT, respectively).

### Cell culture and transfections

Human embryonic kidney cells 293 (HEK293; ATCC) were cultured (Dulbecco’s modified Eagle’s medium high-glucose, 10% fetal bovine serum) at 37°C, 5% CO2, and high humidity. Four days prior to the transfection, the cells were plated in a 10-cm plate at an initial density of 1.3 × 10^6^ cells per plate. Expression plasmids and PEI (product #260008-5; Polysciences) were separately diluted in Opti-MEM1 (product #31985-047; Gibco) and mixtures were incubated at room temperature for 25 minutes to allow polyplex formation prior to its addition to the cell culture. In Mock Conrols, a mock transfection was performed without plasmid but with PEI and Opti-MEM1. The mixture was added dropwise onto the cells. The cells were dissociated 48 hours after transfection by application of Dulbecco’s phosphate-buffered saline without calcium and magnesium (D-PBS) supplemented with 10 mM ethylenediaminetetraacetic acid (EDTA) for 5 min at 37°C. The cells were pelleted and transferred into a 15 mL tube. Pelleting of intact cells was always carried out by centrifugation at 200 × g for 3 min at room temperature.

### Affinity purifications

The cells were washed with 10 mL of warm PBS (37°C) without EDTA, resuspended in 600 μL of ice-cold lysis buffer (50 mM Tris⋅HCl, pH 7.5, 300 mM NaCl, 10% glycerol, 0.2% Triton X-100, and 1% protease inhibitor mixture (P8340; Sigma)), and lysed by sonication^55^. The lysate was cleared by centrifugation at 14,000 g, 4°C, 10 minutes, and the supernatant was saved. Avidin (A9275; Sigma) was added to the cleared lysate to a final concentration of 0.11 mg/mL and incubated at 4°C for 45 minutes. 10 μL of StrepTactin resin (Sepharose High Performance, Cytiva) were added and incubated at 4°C for 2 hours under gentle agitation. The resin was manually washed three times as follows: The resin was brought to the bottom of the tube by centrifugation at 200 g for 45 s, the supernatant was removed, and 1 mL of wash buffer was added (50 mM Tris⋅HCl, pH 7.5, 300 mM NaCl, 10% glycerol).

After the last wash, we strived to remove as much of the supernatant above the resin as possible. We then combined the elution step of the protein from the resin with the denaturation and alkylation steps that are required for mass spectrometry. To that end, we resuspended the resin with 40 μL of denaturation buffer (8 M Urea, 5 mM DTT, 4 mM biotin) for 30 minutes with occasional gentle mixing. We then added iodoacetamide from a 500 mM stock solution to a final concentration of 15 mM. We incubated the alkylation reaction in the dark for an additional 30 minutes with occasional gentle mixing. The resin was allowed to sink to the bottom of the tube, and the supernatant was collected carefully in order not to disturb the resin pellet.

### Cross-linking

Cell pellets expressing either DUX4, RFPL4A or PTOV1, were lysed in PBS lysis buffer (PBS buffer pH 7.4, 0.2% Triton X-100, and 1% protease inhibitor) as described above. After the addition of avidin to the lysates, two lysates were combined according to the complex being studied (i.e. two full cell pellets, each from a 10 cm plate, were used for each cross-linking experiment), and incubated at 4°C for 45 minutes. 10 μL of StrepTactin resin (Sepharose High Performance, Cytiva) were added and incubated at 4°C for 2 hours under gentle agitation. The resin was brought to the bottom of the tube by centrifugation, the supernatant was removed and immediately replaced by 0.5 mL of warm PBS with one of three cross-linking reagents: 2 mM BS^3^ (bis(sulfosuccinimidyl) suberate), 1% Formaldehyde, and 4mM EDC (N-(3-Dimethylaminopropyl)-N′-ethylcarbodiimide hydrochloride) with 8 mM NHS (N-hydroxysuccinimide). The resin was incubated with the cross-linking reagent at 30°C for 30 minutes, and then washed three times with Tris⋅HCl buffer as described above. Because of the concern that the cross-linking would covalently link the Strep tag to the resin, the mass spectrometry preparation protocol was changed to on-bead digestion. To that end, the denaturation and alkylation steps were as described above, but then the resin was not separated from the supernatant and both were transferred for digestion.

### Mass spectrometry

The high-urea sample was diluted by the addition of 360 μL digestion buffer (25 mM Tris⋅HCl, pH 8.0, 10% acetonitrile). For trypsin digestion 1 μg of trypsin (V5111; Promega) was added, and the sample was incubated for 3 hours at 42°C in a shaker. For trypsin-GluC digestion 1 μg of trypsin (V5111; Promega) and 1 μg of GluC (V1651; Promega) were added, and the sample was incubated for 4 hours at 38°C in a shaker. The digest was desalted using Stage Tips, dried in SpeedVac, and resuspended in 0.1% formic acid. The samples were analyzed by a 60-min 0-to-40% acetonitrile gradient on a liquid chromatography system coupled to a Q-Exactive HF mass spectrometer. Between samples an additional acetonitrile gradient was applied to the analytical column in order to reduce the carryover between samples. The analytical column was an EasySpray 25 cm heated to 45°C. The method parameters of the run were as follows: Data-Dependent Acquisition; Full MS resolution 240,000 ; MS1 AGC target 1e6; MS1 Maximum IT 150 ms; Scan range 375 to 1,800; dd-MS/MS resolution 35,000; MS/MS AGC target 2e5; MS2 Maximum IT 150 ms; Loop count Top 12; Isolation window 1.1; Fixed first mass 130; MS2 Minimum AGC target 800; Peptide match - on; Exclude isotope - on; Dynamic exclusion 45 seconds ; HCD energy (NCE): 26; Charge exclusion: unassigned,1,>7 (for proteomics samples), and unassigned,1,2,>7 (for cross-link samples).

### Analysis of mass spectrometry data

Label-free quantification (LFQ) was performed with MaxQuant 1.5 ^56^ using the default parameters. The sequence database comprised all human proteins downloaded from UniProt^57^ augmented with the sequences of the Nucleocapsid protein of SARS-CoV-2. The “proteinGroups.txt” output file was loaded to Perseus^58^. Reverse proteins and contaminations were filtered out, the data transformed to logarithmic scale, and samples grouped according to replicates. Only proteins identified by more than two peptides were considered. For missing LFQ intensities, values were imputed from a normal distribution. The confidence thresholds were determined by a two-sample test with a permutation-based FDR of 0.5% and “s0” (minimal fold change) value of 2. The Volcano plots present the proteins in the “t-test difference” vs. “-Log t-test p-value” coordinate system. The ranking of the proteins by their abundances was according to their average LFQ across the replicates. Cross-links were identified by FindXL as previously described^35^.

### Structural modeling

AlphaFold-Multimer v3 runs were performed using ColabFold with default parameters (without templates), producing five structures per run. The input sequences to the runs were trimmed according to the cross-link data as described in the main text. The cross-links were used again to select the best-fitting model out of the five, based on the shortest combined Cα-Cα distances spanned by the cross-links on the model.

## Supporting information

Supplemental Data Set 1

Supplemental Data Set 2

## Data Availability

The mass spectrometry data have been deposited to the ProteomeXchange Consortium via the PRIDE^59^ partner repository with the dataset identifier PXD045236.

## Acknowledgments

This work was funded by FSHD Global grant #41 and the FSHD Society grant #Winter2021-5912275806

## Supplementary Information

**Figure S1.**
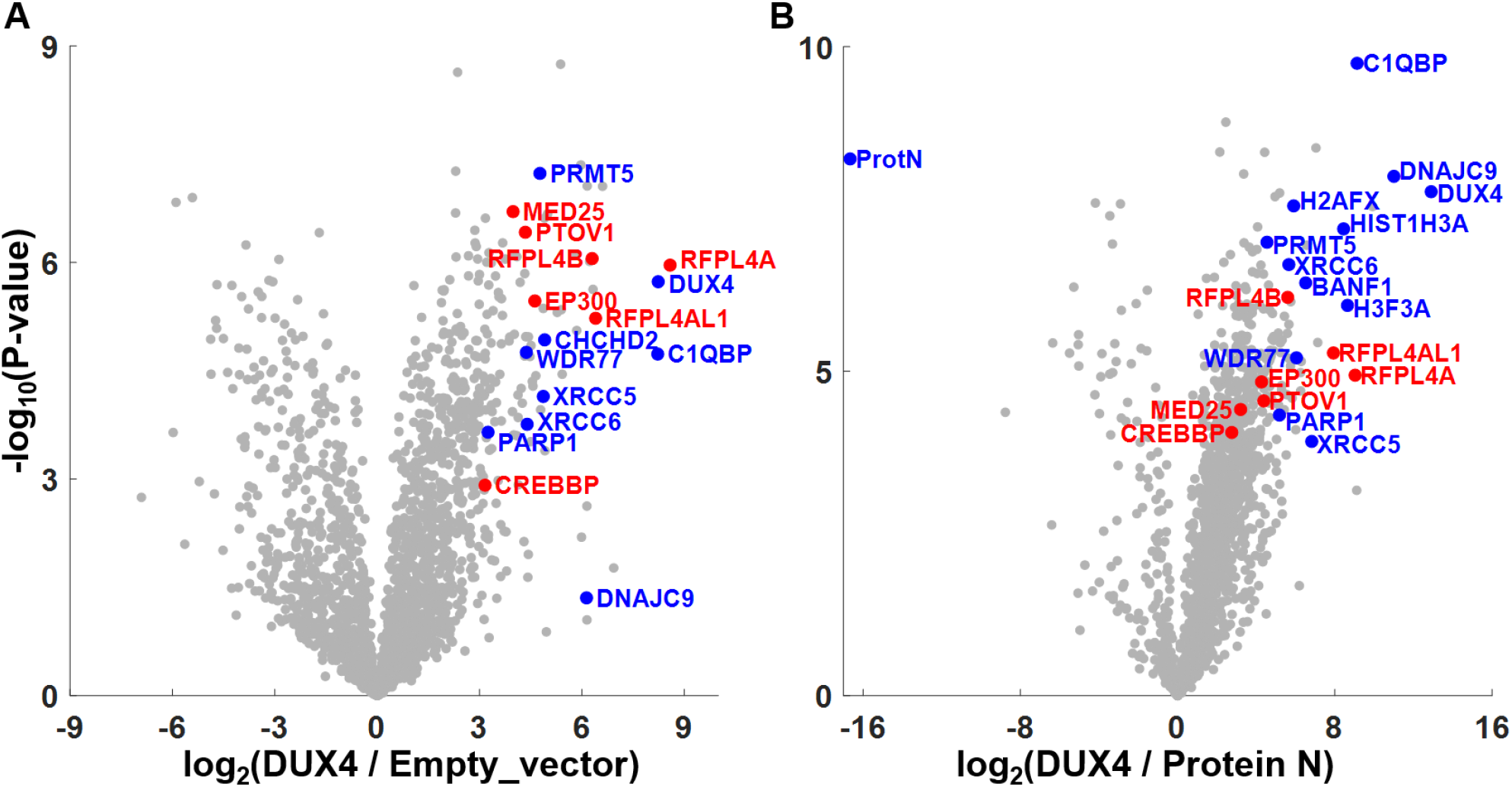
Protein interactions associated with the N-terminal DNA-binding region of DUX4. Results of AP-MS analysis from HEK293 cells that express either DUX4, an empty transfection vector, or the Nucleocapsid protein of SARS-CoV-2 (Protein N). The affinity purification experiments were performed in biological quadruplicates. Proteins that are associated with the N-terminal DNA-binding region and the C-terminal activation region are marked in blue and red, respectively.

**Figure S2.**
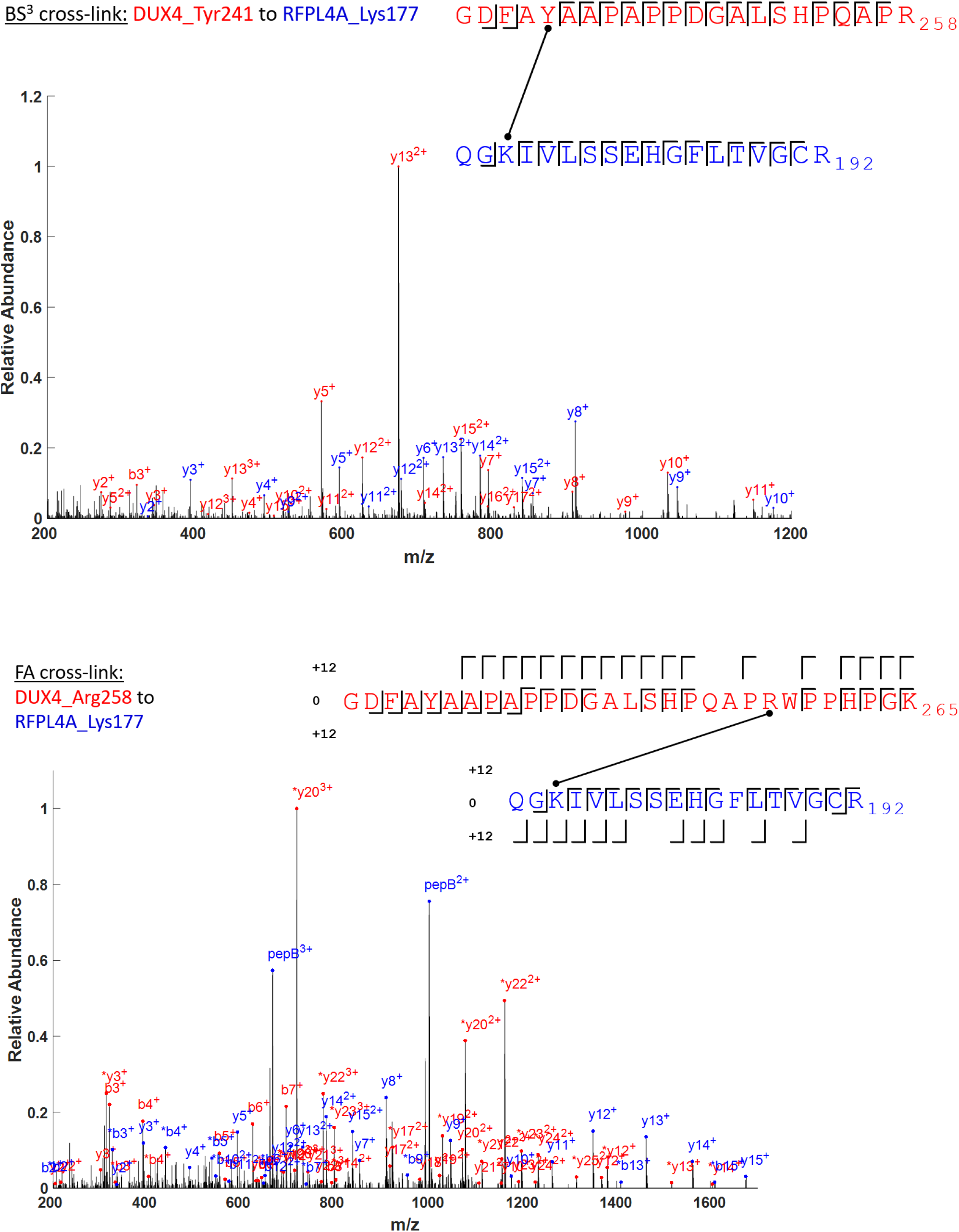
Annotated MS/MS spectra for the DUX4-RFPL4A cross-links. Results for the BS^3^ and Formaldehyde cross-linking reagents are shown in the top and bottom panel, respectively. The annotation of the formaldehyde fragments follow the convention described in Tayri-Wilk et al.^60^.

**Figure S3.**
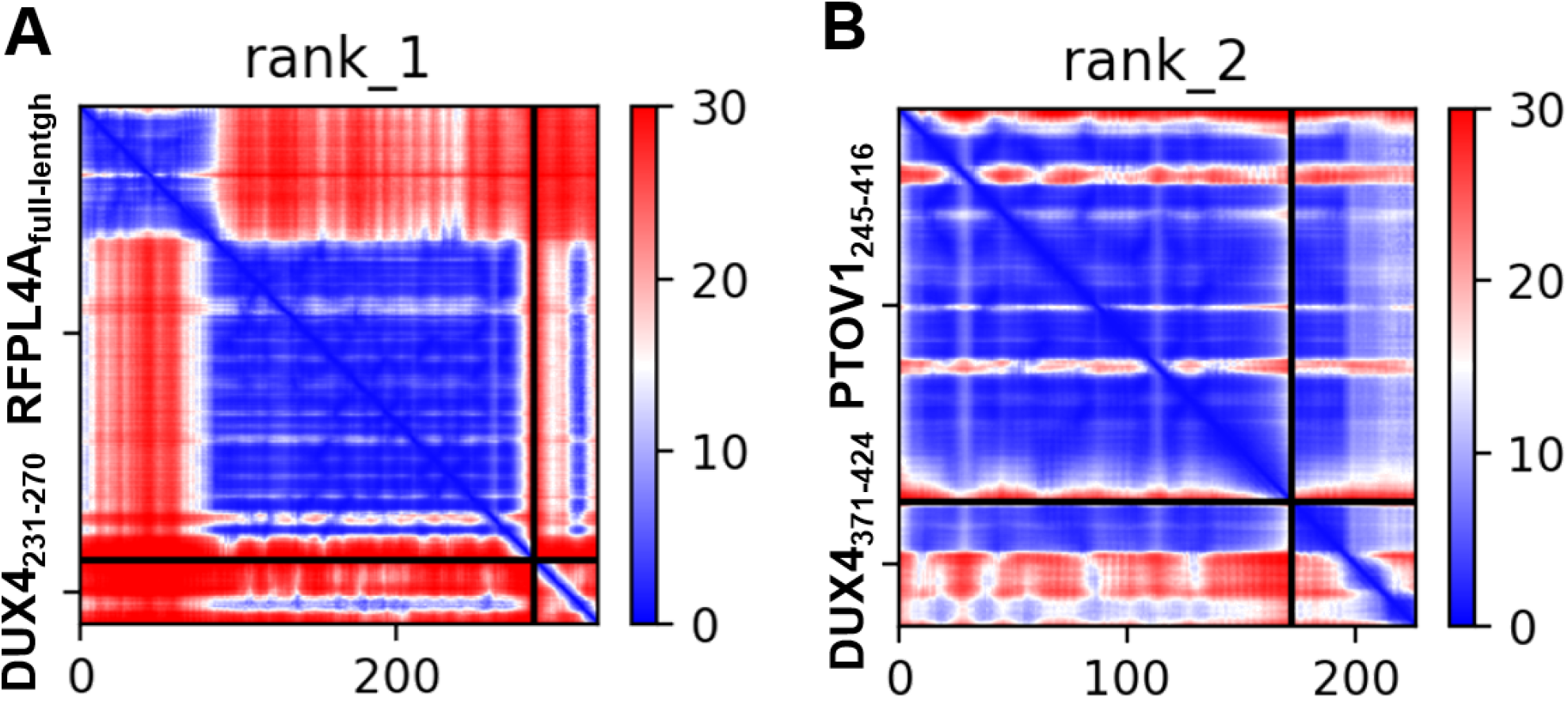
Confidence scores for the AlphaFold Multimer (AFM) structural modeling. **(A)** PAE plot of the AFM run for the selected model of DUX4 and RFPL4A. The pLDDT score was 78.5 and the pTM score was 0.648. **(B)** PAE plot of the AFM run for the selected model of DUX4 and PTOV1. The pLDDT score was 80 and the pTM score was 0.798.

